# Genome-Wide Mining, Characterization and Development of miRNA-SSRs in *Arabidopsis thaliana*

**DOI:** 10.1101/203851

**Authors:** Anuj Kumar, Aditi Chauhan, Mansi Sharma, Sai Kumar Kompelli, Vijay Gahlaut, Johny Ijaq, Krishna Pal Singh, MNV Prasad Gajula, Prashanth Suravajhala, Harindra Singh Balyan, Pushpendra Kumar Gupta

## Abstract

Simple Sequence Repeats (SSRs), also known as microsatellites are short tandem repeats of DNA sequences that are 1-6 bp long. In plants, SSRs serve as a source of important class of molecular markers because of their hypervariabile and co-dominant nature, making them useful both for the genetic studies and marker-assisted breeding. The SSRs are widespread throughout the genome of an organism, so that a large number of SSR datasets are available, most of them from either protein-coding regions or untranslated regions. It is only recently, that their occurrence within microRNAs (miRNA) genes has received attention. As is widely known, miRNA themselves are a class of non-coding RNAs (ncRNAs) with varying length of 19-22 nucleotides (nts), which play an important role in regulating gene expression in plants under different biotic and abiotic stresses. In this communication, we describe the results of a study, where miRNA-SSRs in full length pre-miRNA sequences of *Arabidopsis thaliana* were mined. The sequences were retrieved by annotations available at EnsemblPlants using BatchPrimer3 server with miRNA-SSR flanking primers found to be well distributed. Our analysis shows that miRNA-SSRs are relatively rare in protein-coding regions but abundant in non-coding region. All the observed 147 di-, tri-, tetra-, penta- and hexanucleotide SSRs were located in non-coding regions of all the 5 chromosomes of *A. thaliana*. While we confirm that miRNA-SSRs were commonly spread across the full length pre-miRNAs, we envisage that such studies would allow us to identify newly discovered markers for breeding studies.

## 1. Introduction

## 2. Methodology

### 2.1. Computational identification and discovery of miRNA-SSRs in *A. thaliana* genome

**Figure 1.**
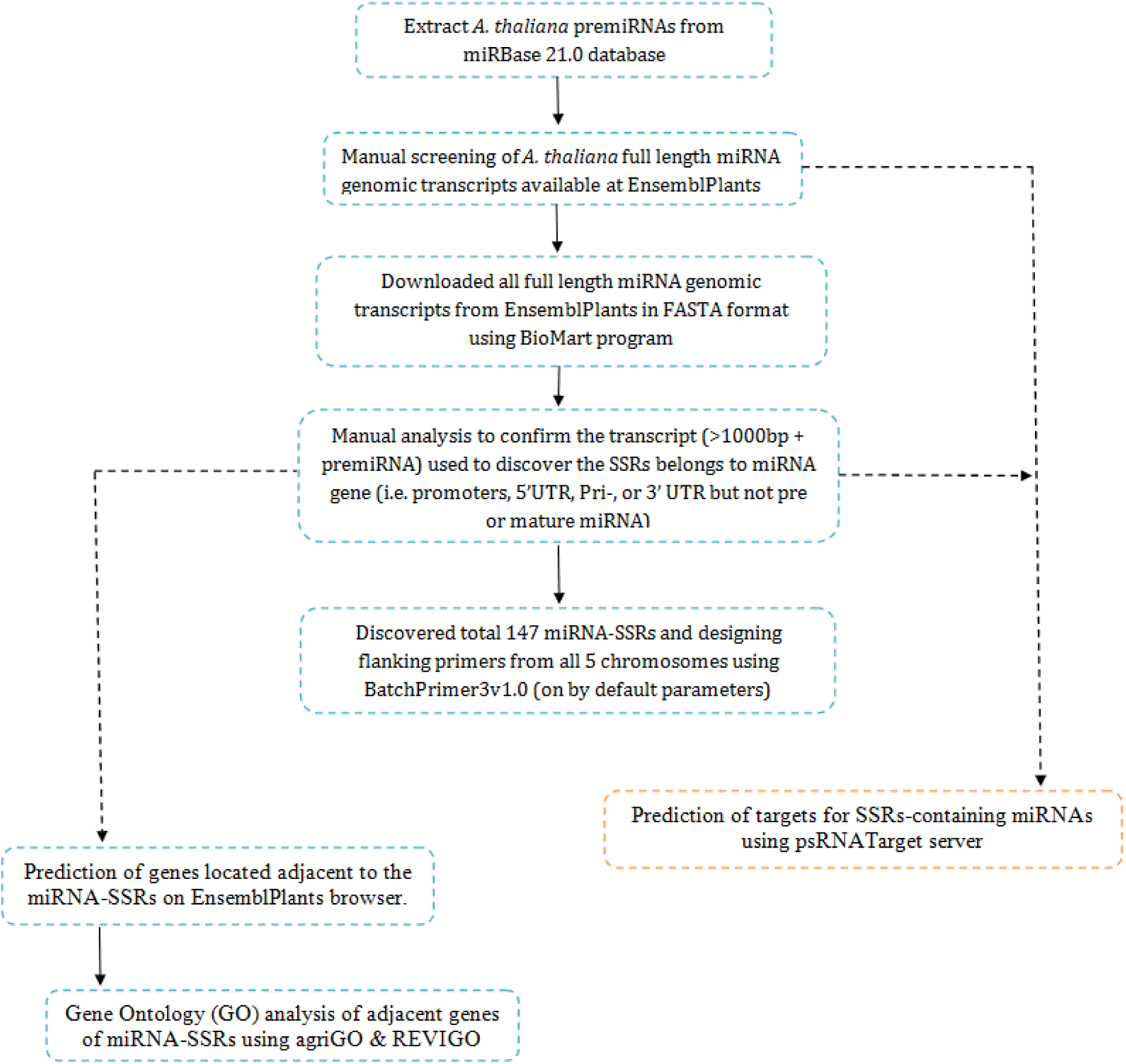
Pipeline used for discovery of miRNA-SSRs in *A. thaliana*.

### 2.2. Computational Prediction of SSRs-containing miRNAs

### 2.3. Prediction of genes adjacent to identified miRNA-SSRs and analysis of enriched gene ontologies (GO)

## 3. Result and Discussion

### 3.1. Dinucleotide repeats were found to outnumber other repeats

In the present study, 147 miRNA-SSRs were discovered among 169 pre-miRNA genomic transcripts of *A. thaliana* genome (Table. 1). We found that dinucleotide SSR repeats (48/147) outnumbered the other repeats; primers designed for 45 of these dinucleotide repeats while no primers were designed for the remaining three SSRs including (AC)_7_ associated with miR164b, (AT)_7_ associated with miR165b and and (TA)_10_ associated with miR832A. Ten (10) different classes of dinucleotide SSR repeats were found in all premiRNA transcripts of *A.thaliana* and the largest count of dinucleotide repeat was TA. (Fig.2). While trinucleotide miRNA-SSR repeats were found to be less than dinucleotide repeats, only one of 38 repeats was found with no SSR flanking primer (TTC with miR837a and SSR length - 12). Nevertheless, there were 37 SSR flanking primers found to be associated with them. Within 15 different classes of trinucleotide miRNA-SSRs repeats, TTC and CTT with same number of counts formed the highest count of trinucleotide repeats (Fig.2)

**Table 1.**
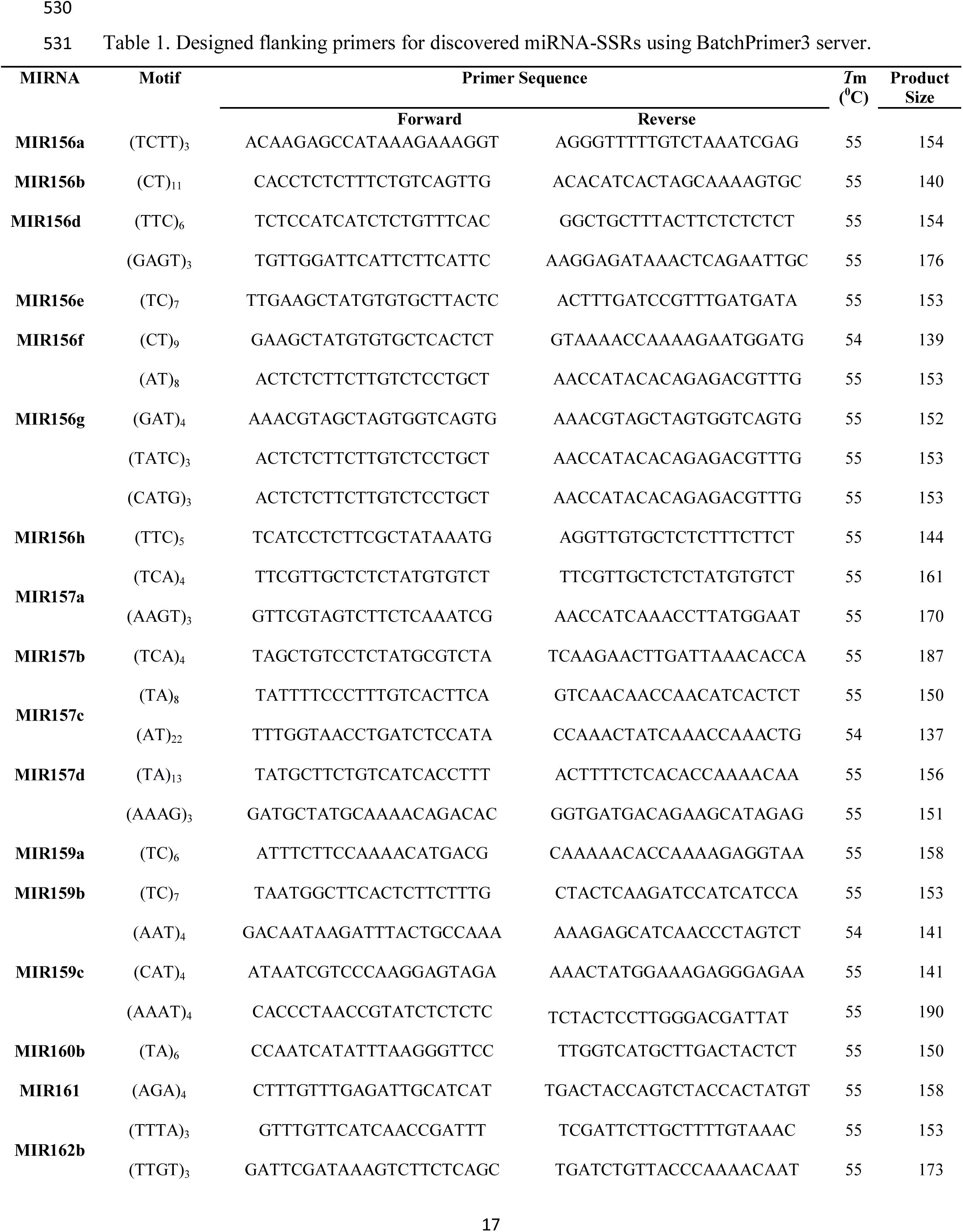
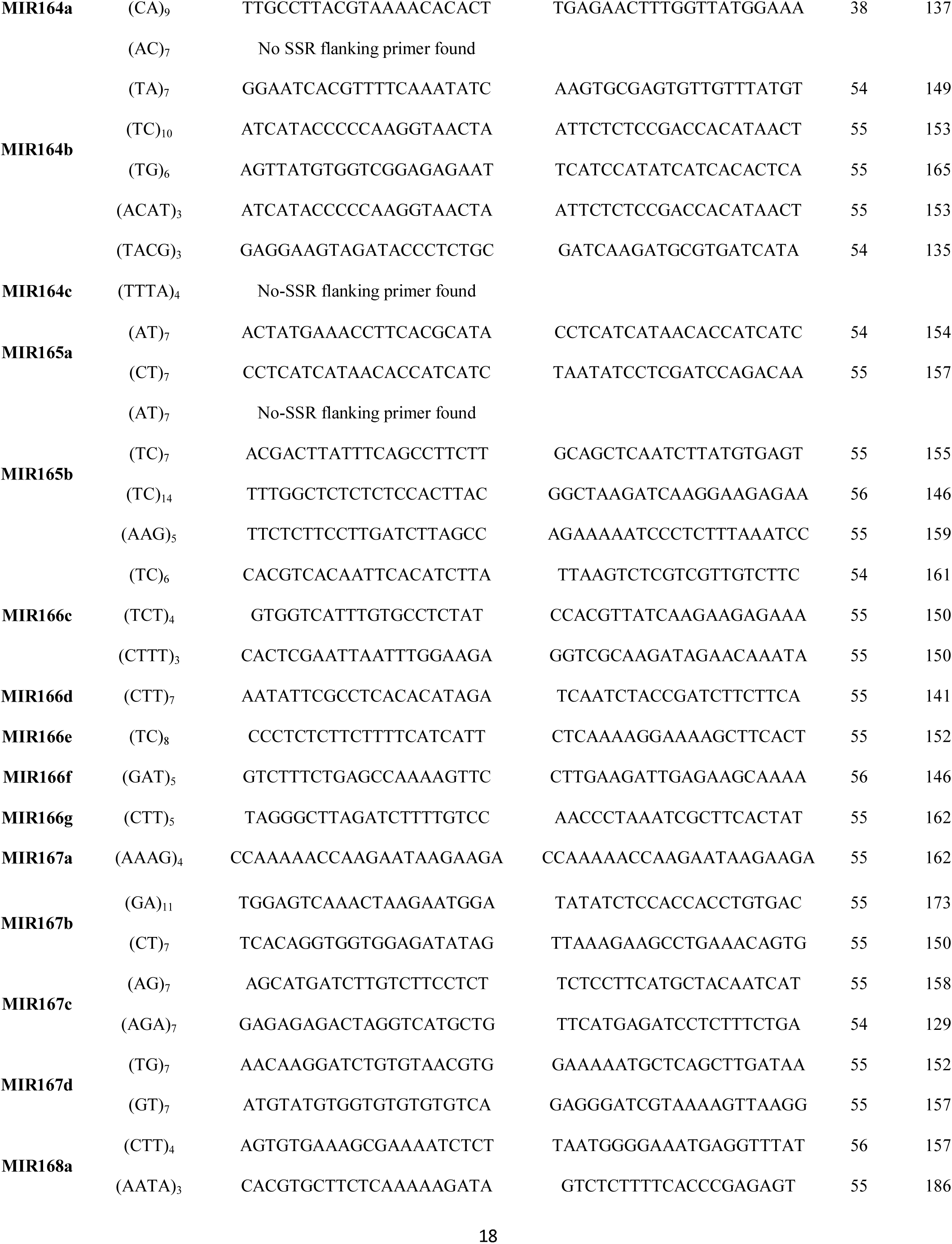
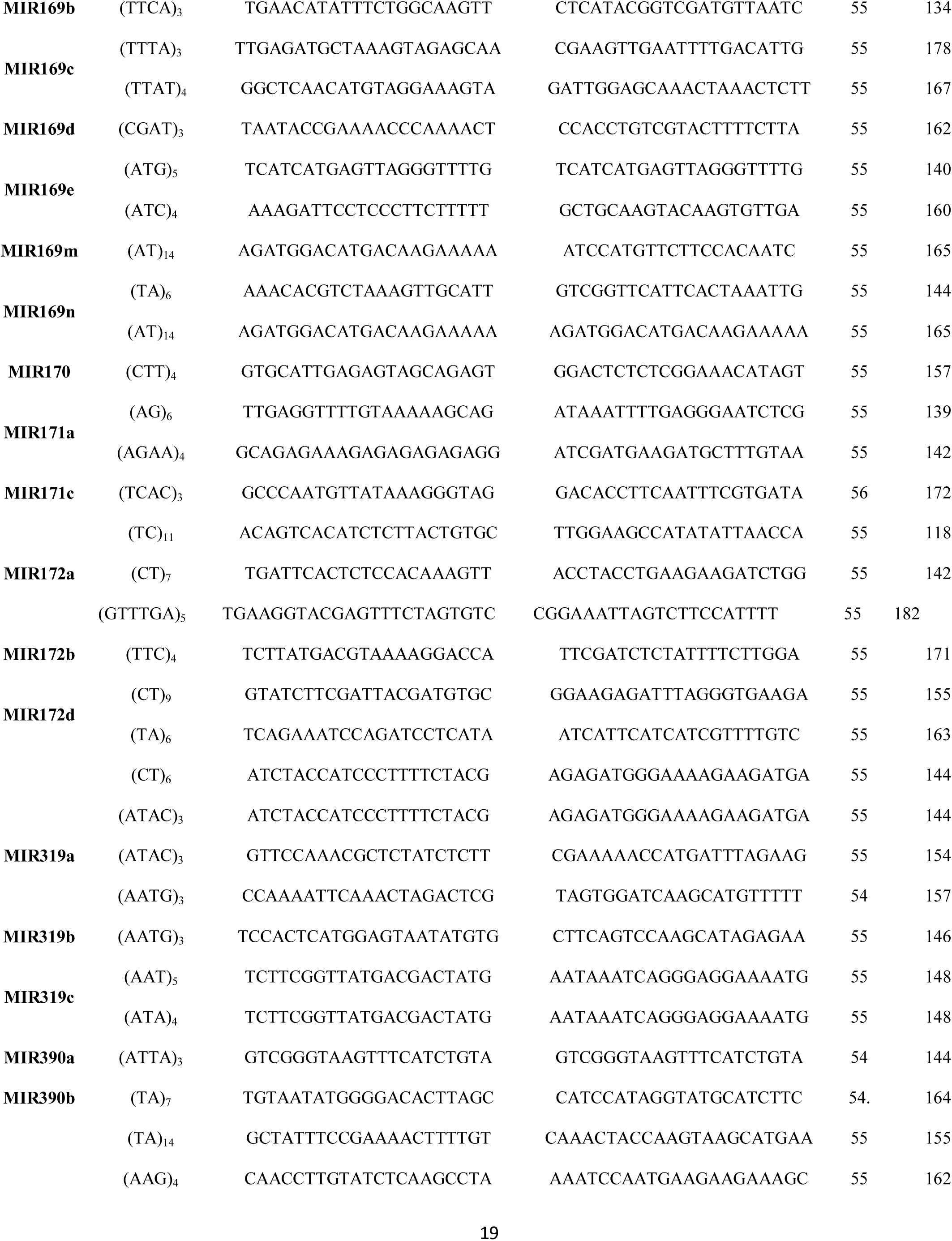
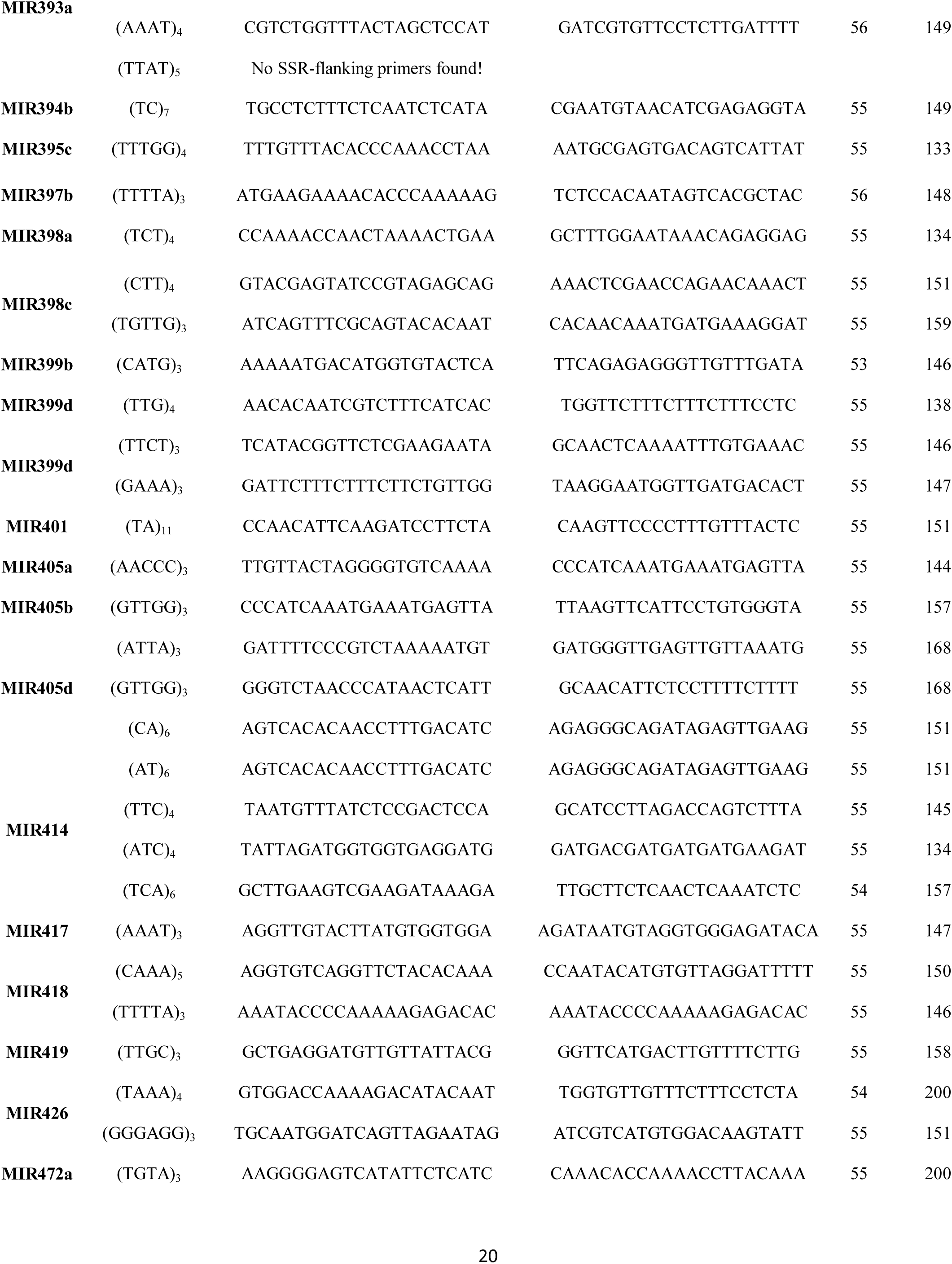
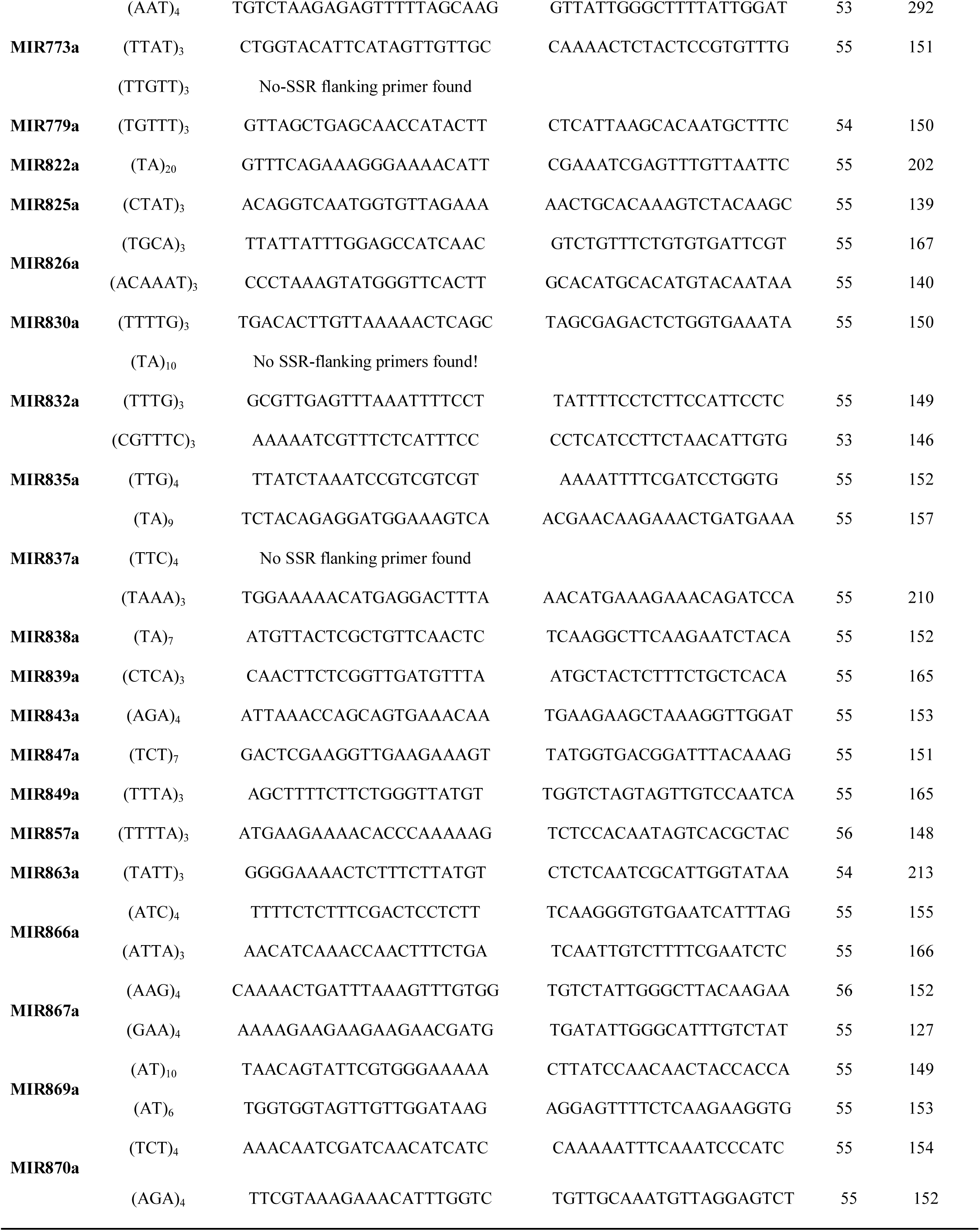
Designed flanking primers for discovered miRNA-SSRs using BatchPrimer3 server.

The tetranucleotide miRNA-SSRs (46) were found to be more than trinucleotide repeats but less than dinucleotide repeats. Primers flanking two SSRs *viz.* (TTTA)_n_, and (TTAT)_n_ for miR164c and miR394a, respectively could not be designed (TTTA)_n_ repeats was most abundant among the tetranucleotide repeats in discovered miRNA-SSRs. (Fig. 2). The pentanucleotide SSRs in pre-miRNA transcripts of *A. thaliana* were least frequent. Out of the 12 of the 147 miRNA-SSRs, were pentanucleotide repeats. Primers flanking to 11 miRNA-SSRs were designed and no primers could be designed for, (TTGTT)_3_ associated with miR777a. Only eight classes of pentanucleotide SSR repeats were found in all pre-miRNA transcripts of *A. thaliana* and TTTTA was found as topmost count of pentanucleotide SSRs (Fig. 2).The hexanucleotide miRNA-SSRs were least common and these belonged to (GTTTGA)_n_, (GGGAGG)_n_, (ACAAAT)_n_, and (CGTTTC)_n_ classes to be associated with flanking primers and remarkably distributed across all 5 chromosome in *A.thaliana* genome (Fig. 3). The chromosomes 1 and 5 have maximum miRNA-SSRs, while chromosome 3 has minimum number of miRNA-SSRs (Fig. 3).

**Figure 2.**
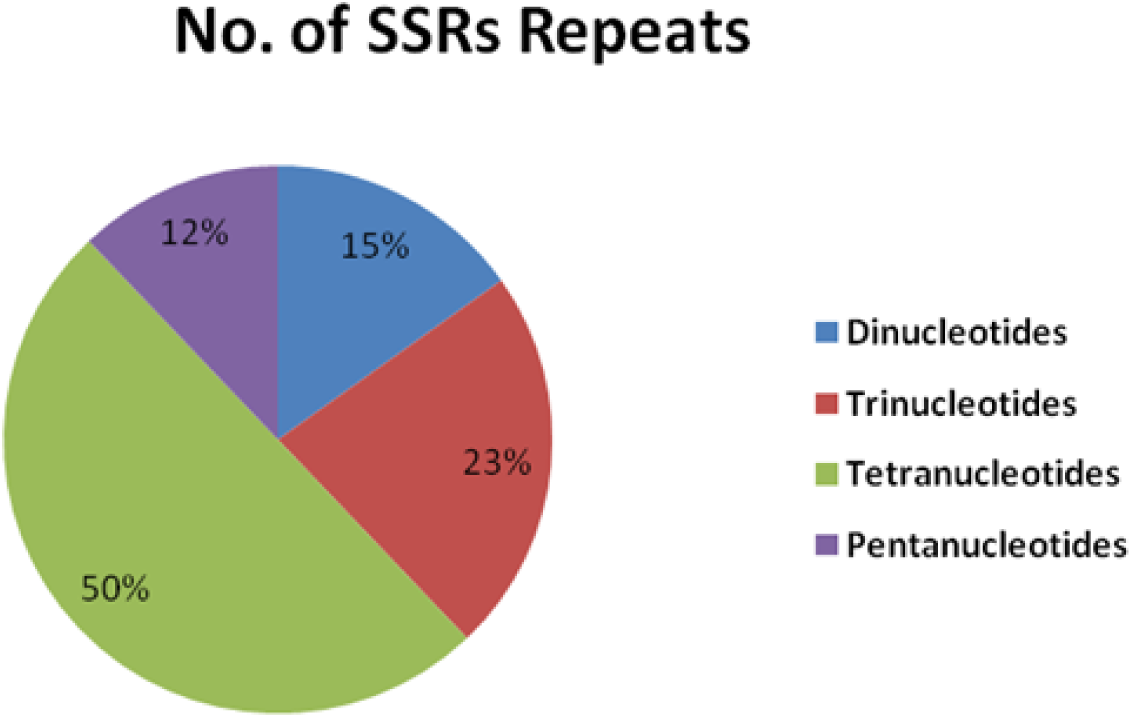
Incidence and number of di, tri, tetra, and pentanucleotide miRNA-SSRs.

**Figure 3.**
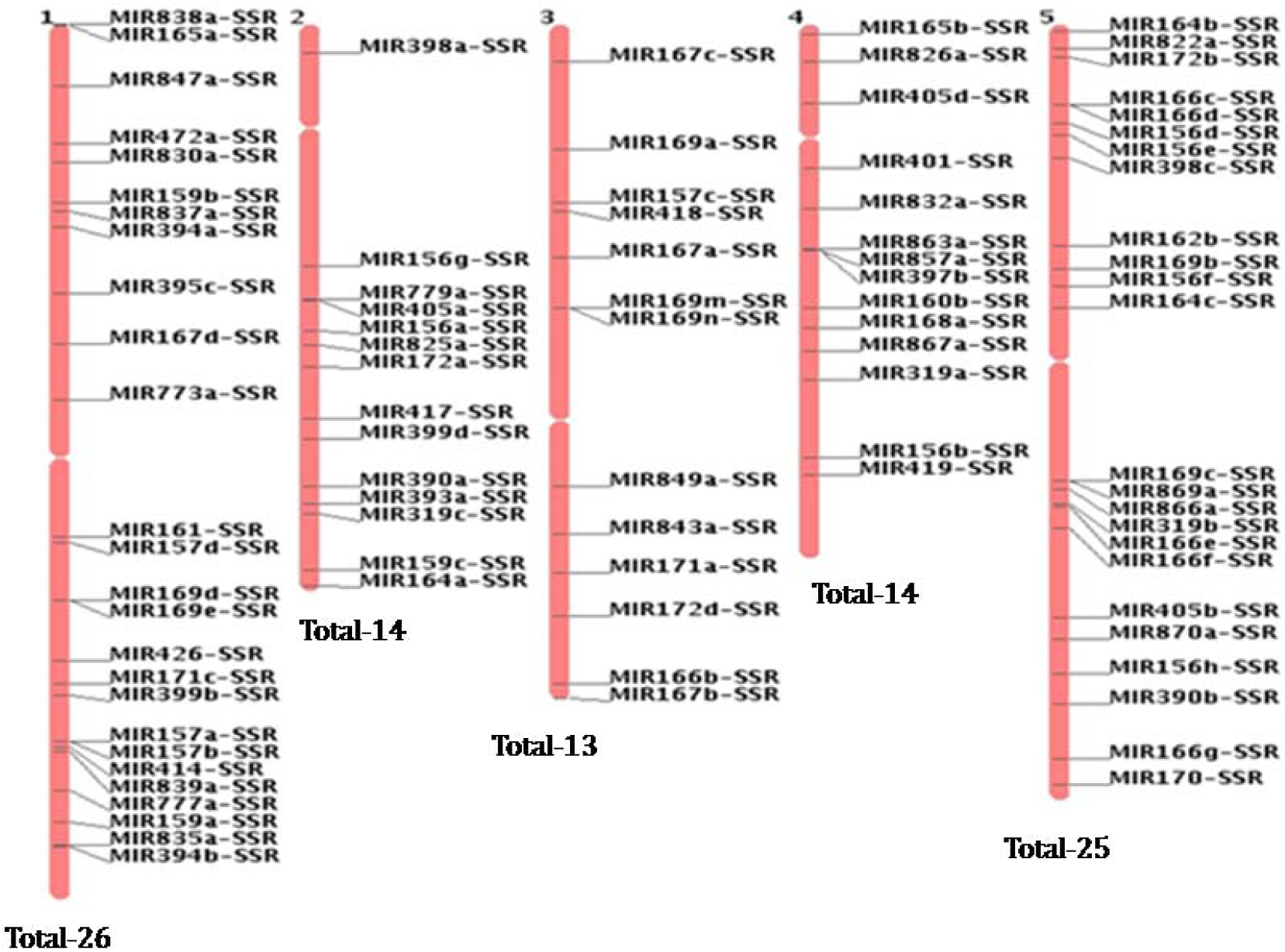
Chromosomal locations of discovered miRNA-SSRs in *A.thaliana* geneome.

### 3.2. Conservation of SSR loci spanning flanking regions

The miRNA-SSR polymorphism will provide trait-related molecular markers at the specific chromosomal loci, which in turn would depend on the number of indels in the flanking regions. Whether or not they are dinucleotide repeats or compound repeats is dependent not only on variances at the each repeat unit of the sequences, but also on how they are arranged or distributed across the genome. As we observed such repeats, it would be interesting to examine their locus specific polymorphism to allow their physically mapping. It would be interesting to see if they can serve as unknown tagged sites which in turn would depend on the presence of a particular sequence tagged region or sequence tagged sites (STS). These STS’ in principle can be used as potential markers.

### 3.3. SSRs-containing miRNAs targeted diverse set of TFs

On the basis of the biogenesis of miRNAs in plants, a homology search-based method was used to predict the targets for SSRs-containing miRNA in *A. thaliana* using psRNATarget. The SSR-containing miRNAs were used as queries to predict potential mRNA targets in the Arabidopsis genome annotation (TAIR10). This search revealed that 90 SSR-containing miRNAs identified 698 target genes, with each SSR-containing miRNA predicting more than one gene (Table S1). Most of the SSR-containing miRNAs targeted a number of TFs families including WRKY, MADS, MYB, NAC, bHLH, AP2/EREBP, ARF etc., which play an important role in different metabolic and regulatory processes such as stress response, transcriptional regulation, signal transduction, growth, development, nutrient uptake, nutrient transport and nutrient assimilation (Table 2). The values of UPE for targeted gene ranged from 3.238 to 24.941.

**Table 2.**
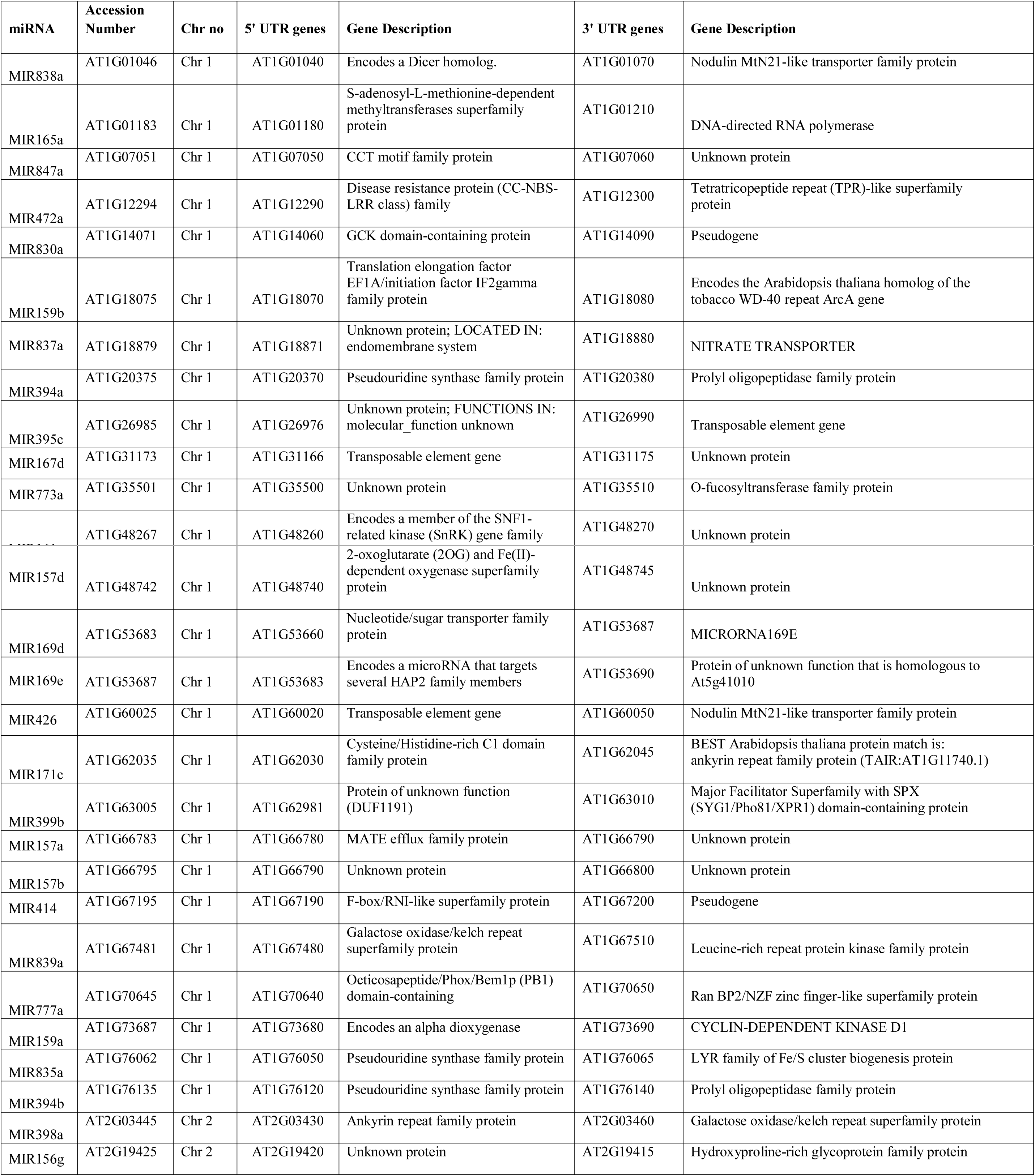
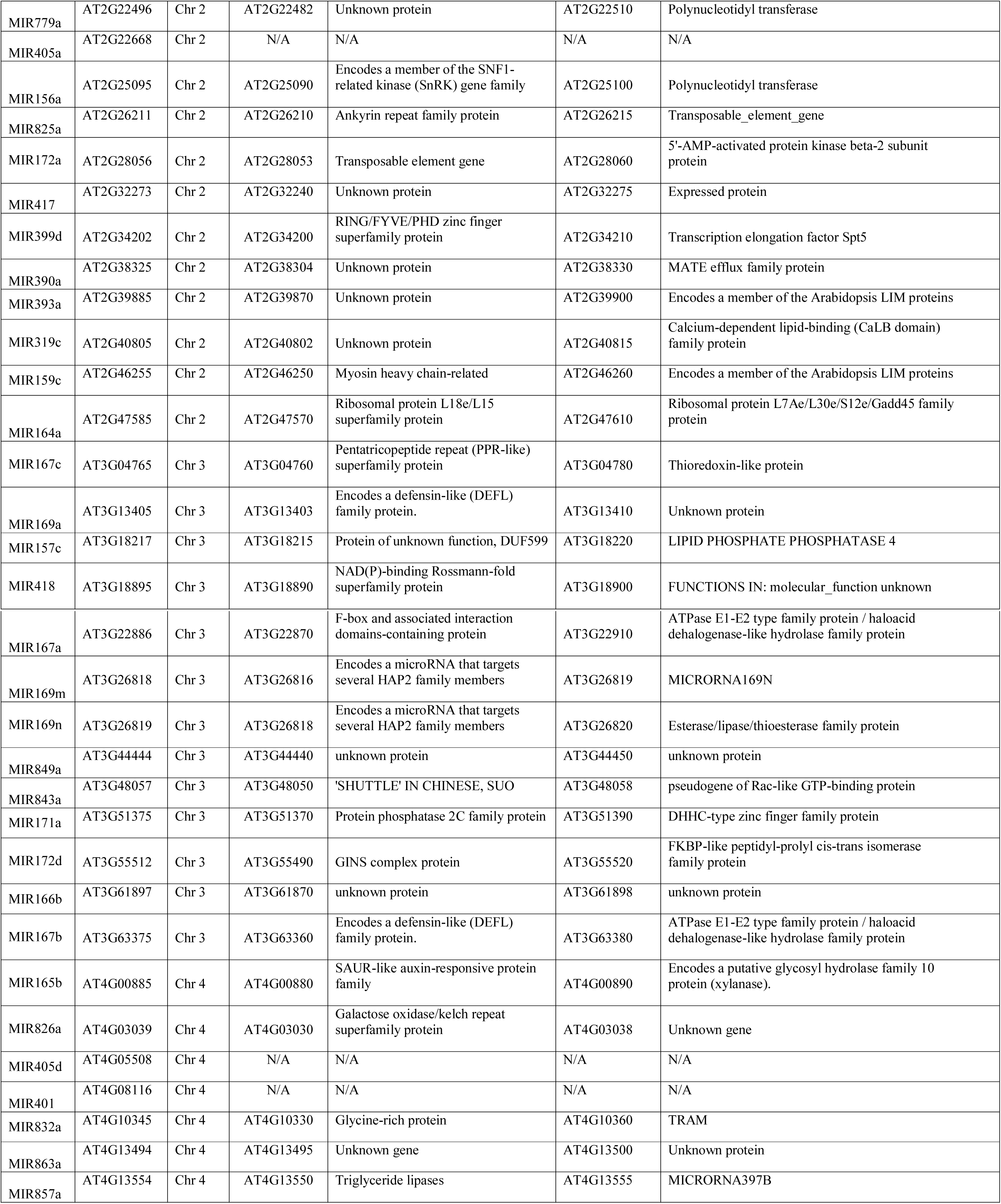
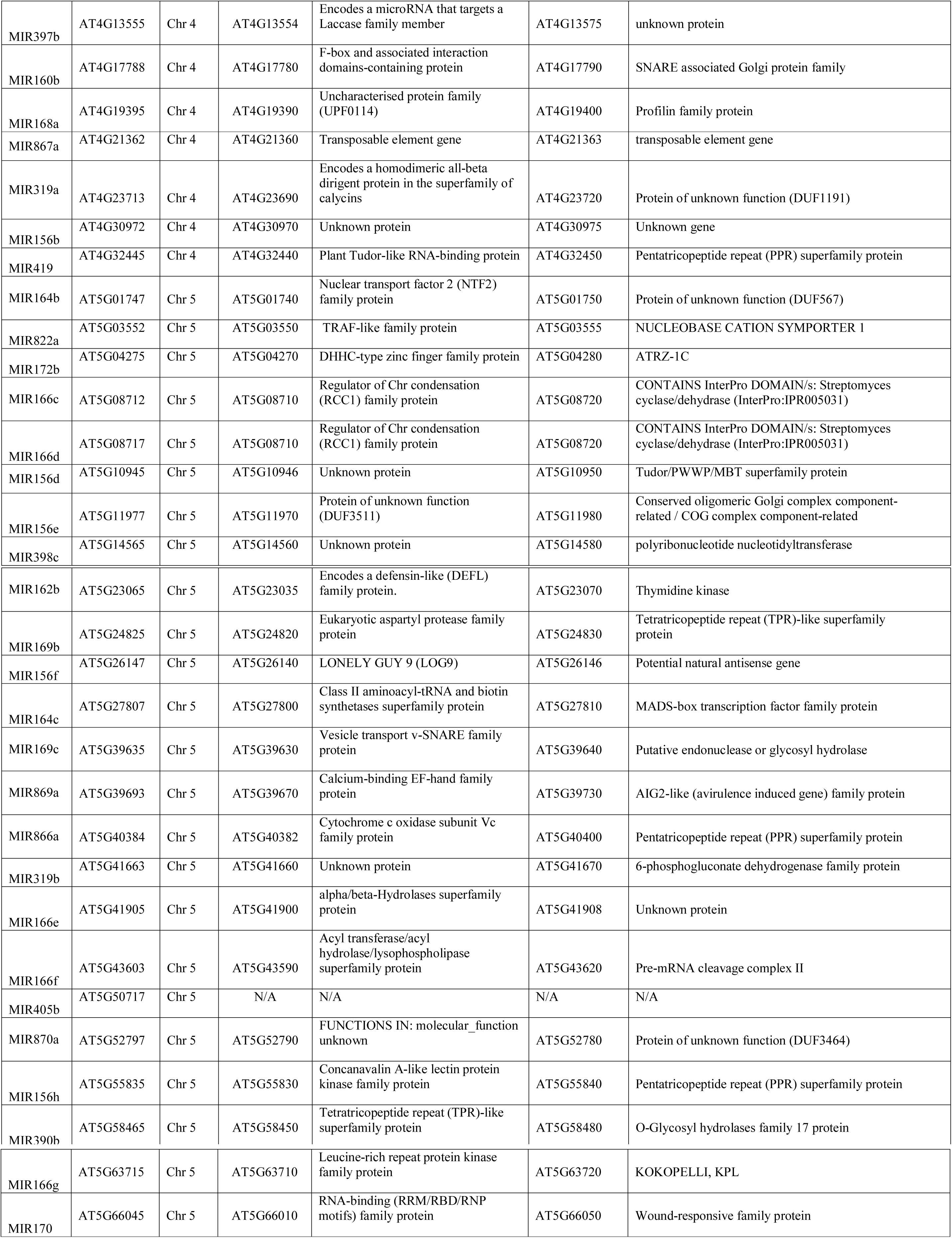
Genes located adjacent to the miRNA-SSRs.

### 3.4. Prediction of genes adjacent to identified miRNA-SSRs and GO analysis

In order to predict the genes adjacent to SSR containing miRNAs, representing 5′ UTR and 3′ UTR sites TAIR 9 was manually curated. Based on length and chromosomal location, a diverse set of adjacent genes were predicted both in n5′ UTR and 3′ UTR regions (Table. 2). Predicted adjacent transcripts revealed that SSR containing miRNAs are associated with different genes in network form, which play a pivotal role in gene regulation. However effect of miR-SSR on adjacent genes and vice-versa need to be studied in detail.

To evaluate the biological significance of the adjacent genes to SSR containing miRNAs in Arabidopsis it is important to have the gene ontology (GO) descriptions i.e., detailed annotations of gene function, biological process it is involved, and cellular location of the gene product. The potential functions were predicted by searching against GO database using agriGO and REVIGO server. Predicted adjacent transcripts were subjected to singular enrichment analysis (SEA) embedded in agriGO to identify enriched GOs. SEA designed to identify enriched GO terms in a list form of microarray probe sets or gene identifiers available in database. Finding different enriched GO terms corresponds to finding enriched biological facts, and term enrichment level was judged by comparing query list to a background population from which the query list is derived. In this study the background query list comprised of 27,416 protein coding genes from the updated TAIR (https://www.arabidopsis.org/index.jsp). Fig. 4 wholly reflects the categorization of adjacent genes based on biological process, cellular component and molecular function. Adjacent genes were divided into 14 GO categories. Among the adjacent gene transcripts, GOs associated with response to stimulus, cellular biosynthetic process, nitrogen compound metabolic process, nucleobase, nucleoside, cellular macromolecule metabolic process, protein metabolic process, transport activity, RNA metabolic process, gene regulation and binding (Fig.5).

**Figure 4.**
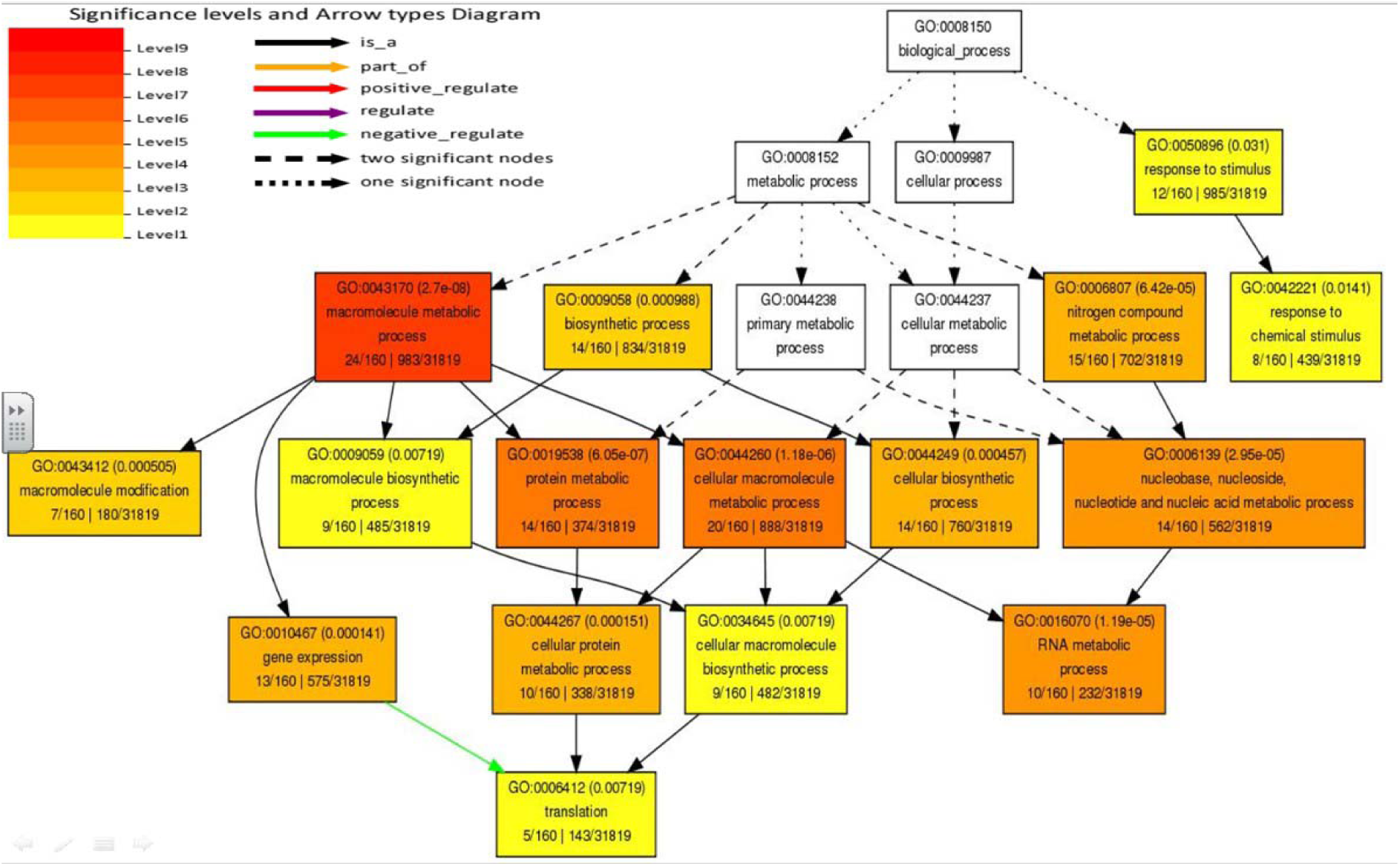
GO classifications of adjacent genes to SSR containing miRNAs.

**Figure 5.**
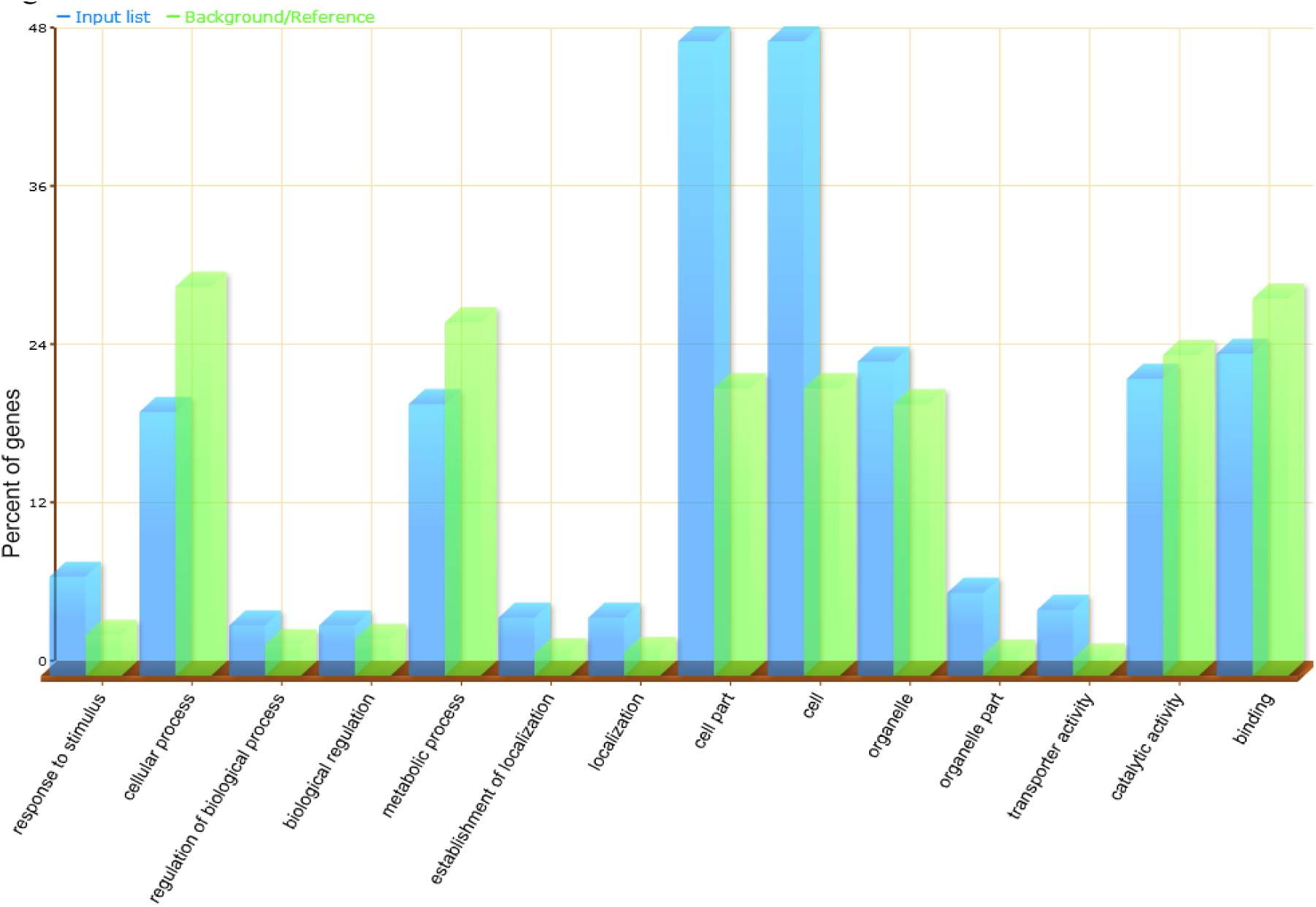
GO analysis of adjacent genes to SSR containing miRNAs: box reflects the GO term number, the p-value in parenthesis, and GO term. The first pair of numerals shows the number of adjacent genes in the input list associated with that GO term and the number of genes in the input list. The second pair of numerals represents the number of genes associated with the particular GO term in the TAIR database and the total number of Arabidopsis genes with GO annotations in the TAIR database. The box colours indicates levels of statistical significance with yellow = 0.05; orange = e-05 and red = e-09.

In order to reduce the number of GO terms, enriched GO categories with false discovery rates (FDR) < 0.05 from AgriGO analysis were submitted to the REVIGO (REduce and Visualize GO) server. Using the Uniprot (http://www.uniprot.org/) as background and the default semantic similarity measure (Simrel), this analysis clearly showed that biological processes associated with metabolism, localization, nitrogen regulation, regulation of transcription were significantly overrepresented among the adjacent genes to SSR containing miRNAs in Arabidopsis (Fig.6).

**Figure 6.**
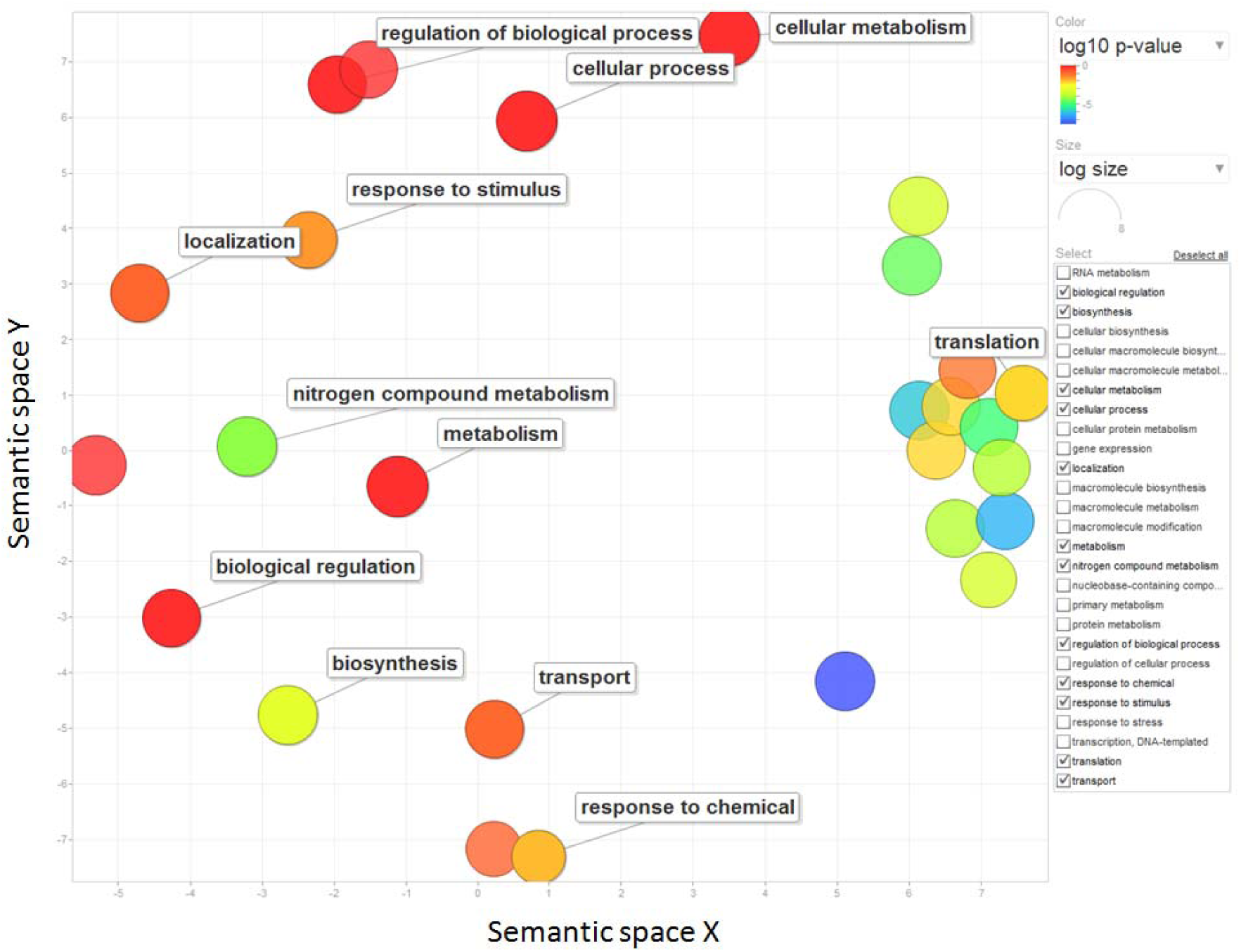
GO analysis of adjacent genes to SSR containing miRNAs using REVIGO: The scatter plot represents the cluster representatives (terms remaining after reducing redundancy) in a two-dimensional space derived by applying multi-dimensional scaling to a matrix of GO terms semantic similarities. Bubble color indicates the p-value for the false discovery rates derived from the AgriGO analysis. The circle size represents the frequency of the GO term in the uniprot database (more general terms are represented by larger size bubbles).

### 3.5. Taking an analogy with long non-coding RNAs

## 4. Conclusion

In the present study, we discovered total 147 miRNA-SSRs from 169 pre-miRNAs representing full length genomic transcripts of *A. thaliana*. Our result shows that all the di-, tri-, tetra-, penta and hexanucleotide SSRs were located in non-coding repertoire of all the 5 chromosomes of *A.thaliana* (Fig. 3). While dinucleotide miRNA-SSRs were found to be higher, hexanucleotide miRNA-SSRs were found to be lowest repeats in the pre-miRNA transcripts. It was observed that miRNA-SSRs flanking primers were larger in number for discovered miRNA-SSRs. We firmly consider these candidates could be extended for experimentation for allelic variation. It is important to know that these miRNA-SSRs serve as a source of highly informative molecular markers and aids as a reference for marker assisted breeding in plants. We hope this first report on genome-wide identification and characterization of miRNA-SSRs in *A. thaliana* could serve as a reference for identifying more sequences from non-coding repertoire of the genomes.

## Acknowledgments

AK would like to give his sincere thanks to Mr. Deepak Kumar, Secretary, IT, ST & BT Government of Uttarakhand for encouragement, suggestions and timely help. PKG was awarded a National Academy of Sciences India (NASI) Senior Scientist Platinum Jubilee Fellowship, and INSA Senior Scientist positions during the tenure of which this study was conducted; VG was awarded a Junior Research Fellowship under the same program, and was later awarded the position of SRF/ RA under a DBT project.

## Authors Contributions

AK, AC, SKK, and VG performed the data analysis; KPS and MNVPG manually crosschecked the annotation. KPS assisted AK and AC for preparing the first draft. PS, HSB and PKG conceived, supervised, edited, and finalized the manuscript.

